# A three-sample test for introgression

**DOI:** 10.1101/594333

**Authors:** Matthew W. Hahn, Mark S. Hibbins

**Affiliations:** Department of Biology, Indiana University; Department of Computer Science, Indiana University

**Keywords:** hybridization, ABBA-BABA, Patterson’s D, gene flow, admixture

## Abstract

Many methods exist for detecting introgression between non-sister species, but the most commonly used require either a single sequence from four or more taxa or multiple sequences from each of three taxa. Here we present a test for introgression that uses only a single sequence from three taxa. This test, denoted *D*_3_, uses similar logic as the standard *D*-test for introgression, but by using pairwise distances instead of site patterns it is able to detect the same signal of introgression with fewer species. We use simulations to show that *D*_3_ has statistical power almost equal to *D*, demonstrating its use on a dataset of wild bananas (*Musa*). The new test is easy to apply and easy to interpret, and should find wide use among currently available datasets.

## Introduction

Genome-scale data have revealed extensive evidence for post-speciation introgression across the tree of life (reviewed in Mallet et al. 2016). Many of these analyses have been carried out in a phylogenetic context, using only a single sample from each population or species. Some methods use gene tree topologies themselves as input (e.g. Huson et al. 2005; Meng and Kubatko 2009; Yu et al. 2011; Edelman et al. 2018), while others use counts of shared derived alleles that reflect the underlying topologies (e.g. Green et al. 2010; Lohse and Frantz 2014; Pease and Hahn 2015).

All of these methods depend on the expectation under incomplete lineage sorting (ILS) that the two less-frequent topologies in a rooted triplet should be equal in frequency. Asymmetry in gene tree topologies is taken as evidence for introgression, though ancestral population structure can produce similar patterns (Slatkin and Pollack 2008; Durand et al. 2011; Lohse and Frantz 2014). Importantly, the need to distinguish among topologies or between ancestral and derived sites using these methods means that at least four taxa must be sampled, and sometimes more (e.g. Pease and Hahn 2015; Elworth et al. 2018).

Here, we present a test for introgression that only requires a single sample from each of three taxa. With three taxa we cannot infer the frequencies of alternative gene tree topologies. Instead, our test is based on a related prediction of the ILS model: that there is also an expected symmetry in the branch lengths among topologies. Using these branch lengths, we can develop a test statistic based on pairwise distances to detect the presence of introgression.

## New Approaches

### A test for introgression

Assume that lineages *A* and *B* are sister in the species tree, with divergence time *t*_1_ (measured in units of 2*N* generations), and that the ancestor of *A* and *B* split from lineage *C* at time *t*_2_ (Figure 1a). We refer to gene trees having this topology as *AB*, such that the two discordant topologies are *AC* and *BC* (Figures 1b and 1c, respectively).

**FIG. 1.**
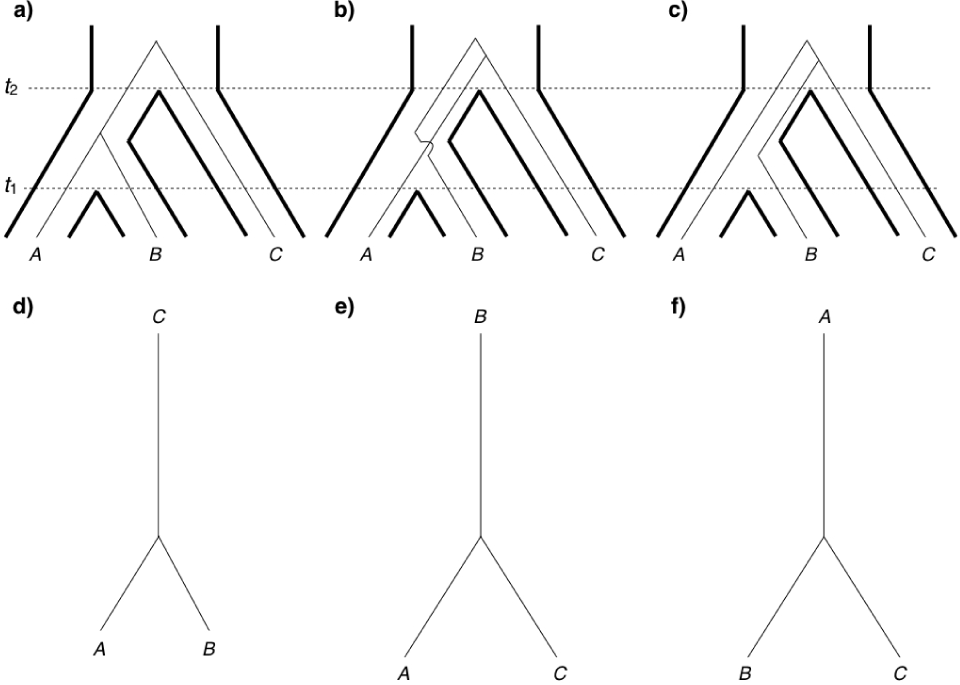
Topologies produced by incomplete lineage sorting. The top row shows the same species tree (thick lines, with divergence times denoted by *t*_1_ and *t*_2_) within which three different topologies arise: a) *AB*1, b) *AC*, and c) *BC*. The bottom row shows the same unrooted topologies as in a-c, with approximate branch lengths.

When ILS is the only cause of gene tree incongruence, topology *AB* may be generated in two different ways, with different expected frequencies and branch lengths. Looking backwards in time, we refer to the topology in which lineages *A* and *B* coalesce before *t*_2_ as *AB*1 (this is the history shown in Figure 1a). Alternatively, the same topology can occur when these lineages coalesce in the ancestral population of all three lineages; we refer to this topology as *AB*2.

The expected frequencies of these four topologies are (Hudson 1983):

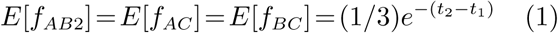

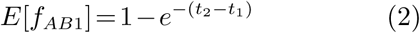

As mentioned in the Introduction, here we see that the two discordant topologies (*AC* and *BC*) are expected to have the same frequencies.

The same model leads naturally to expectations for the times to coalescence between lineages in each of the different topologies. Here we focus on the expected times to coalescence between *B* and *C* (*t*_*B−C*_) and between *A* and *C* (*t*_*A−C*_). These times are (Hibbins and Hahn 2019):

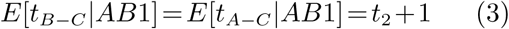

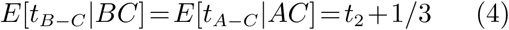

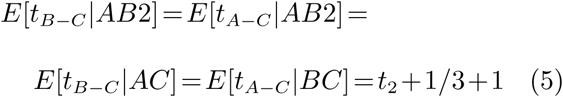

These times can be transformed into genetic distances between tip sequences by assuming an infinite sites mutation model and multiplying by two to account for mutations along both lineages since their common ancestor. Summing the weighted length of branches between any two taxa across all possible topologies leads to the following expected distances:

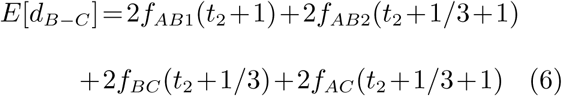

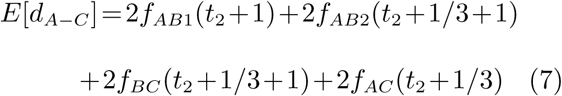

(leaving off the shared mutation parameter, *µ*, for clarity). Due to the underlying symmetries in topology frequencies and branch lengths under ILS, the expected values of *d*_*B−C*_ and *d*_*A−C*_ are exactly the same. Notably, these expectations hold for distances calculated without rooted gene trees or polarized substitutions (e.g. Figure 1d-f).

Given these results, a natural test of the ILS-only model can be formed using the statistic:

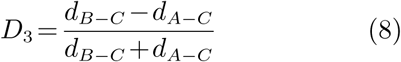

Because the two terms in the numerator have the same expected values under ILS alone, the expectation of *D*_3_ is 0. The denominator is a normalizing factor that bounds *D*_3_ between −1 and +1.

*D*_3_ can be significantly different from zero in the presence of gene flow. While the exchange of alleles between lineages *A* and *B* will have no effect on *D*_3_, unequal amounts of introgression between either *B* and *C* (Figure 2a) or *A* and *C* (Figure 2b) can lead to deviations from zero. This occurs because gene flow between a pair of non-sister lineages leads to a breakdown in the symmetry of branch lengths predicted under ILS alone. In particular, introgression between *B* and *C* leads to both more trees with a *BC* topology and a shorter pairwise distance between these two lineages (Figure 2a). As a result, *d*_*B−C*_ will be smaller than *d*_*A−C*_, leading to a negative value of *D*_3_. Conversely, gene flow between *A* and *C* leads to positive values of *D*_3_. Exact expectations for *D*_3_ in the presence of introgression are presented in the Appendix.

**FIG. 2.**
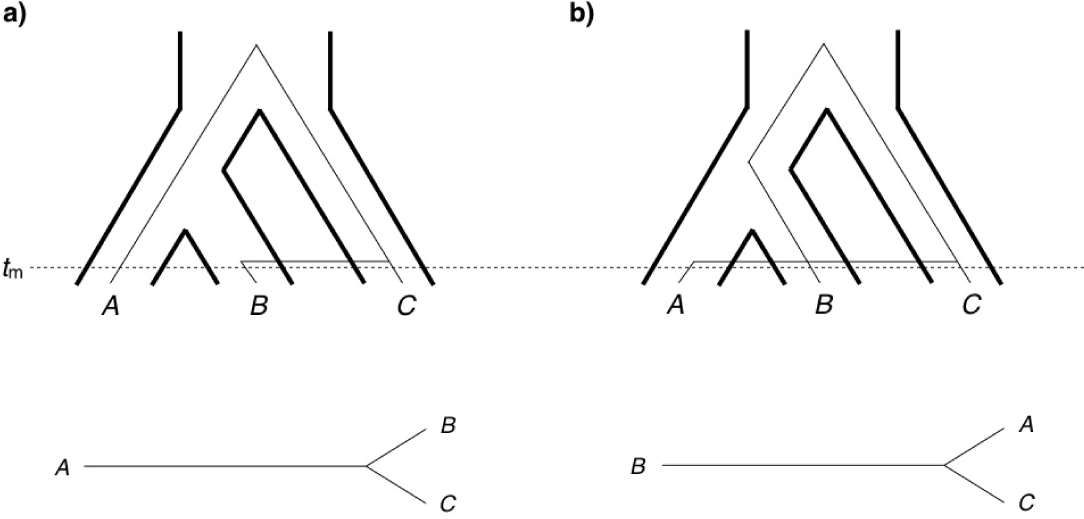
Topologies produced by introgression. The top row shows the same species tree as in Figure 1, but with introgression between a) lineages *B* and *C*, or b) lineages *A* and *C*. Introgression occurs at time *t*_*m*_ in both scenarios. The bottom row again shows the approximate unrooted topologies resulting from introgression. Note how the distance between lineages a) *B* and *C*, or b) *A* and *C* are smaller than in the ILS-only case (Figure 1).

## Results and Discussion

### Application of *D*_3_

The *D*_3_ test is straightforward to carry out, requiring only pairwise distances between three species. Ideally, distances should be calculated from regions for which all three lineages have sequences present in the alignment. This will avoid biases that could possibly occur if regions with different ancestral effective population sizes (for example, in regions with different recombination rates; Pease and Hahn 2013) are sampled unequally for the two relevant distances. Otherwise, variation in either *N* or *µ* across sites should not affect the expectation of *D*_3_.

As an example application of this method, we calculated *D*_3_ for whole-genome data from three sub-species within *Musa acuminata* (wild bananas; the alignment can be found at https://doi.org/10.6084/m9.figshare.7924727.v1). As was found using the original *D*-test on these three taxa and an outgroup (Rouard et al. 2018), *D*_3_ indicated gene flow between the subspecies *M. a. malaccensis* and *M. a. burmannica* (*D*_3_=0.06; *P* <0.0001). The significance of *D*_3_ was determined by a block bootstrap of the Musa alignment, as is normally done for the *D*-test (Green et al. 2010).

### Statistical power of *D*_3_ and comparison with *D*

We tested the power of *D*_3_ to detect gene flow with increasing levels of introgression (Figure 3a). As the fraction of the genome introgressed approaches 10%, *D*_3_ can detect gene flow in 94% of simulated datasets (at *P* < 0.05). This demonstrates that *D*_3_ has good power to detect introgression. In contrast, when there is no gene flow (*γ*=0), the proportion of false positives is the number we would expect at this significance threshold (Figure 3a). We can also see that the expected values of *D*_3_ under different levels of introgression (calculated according to the equations given in the Appendix) closely match the mean of simulated datasets (Figure 3a).

**FIG. 3.**
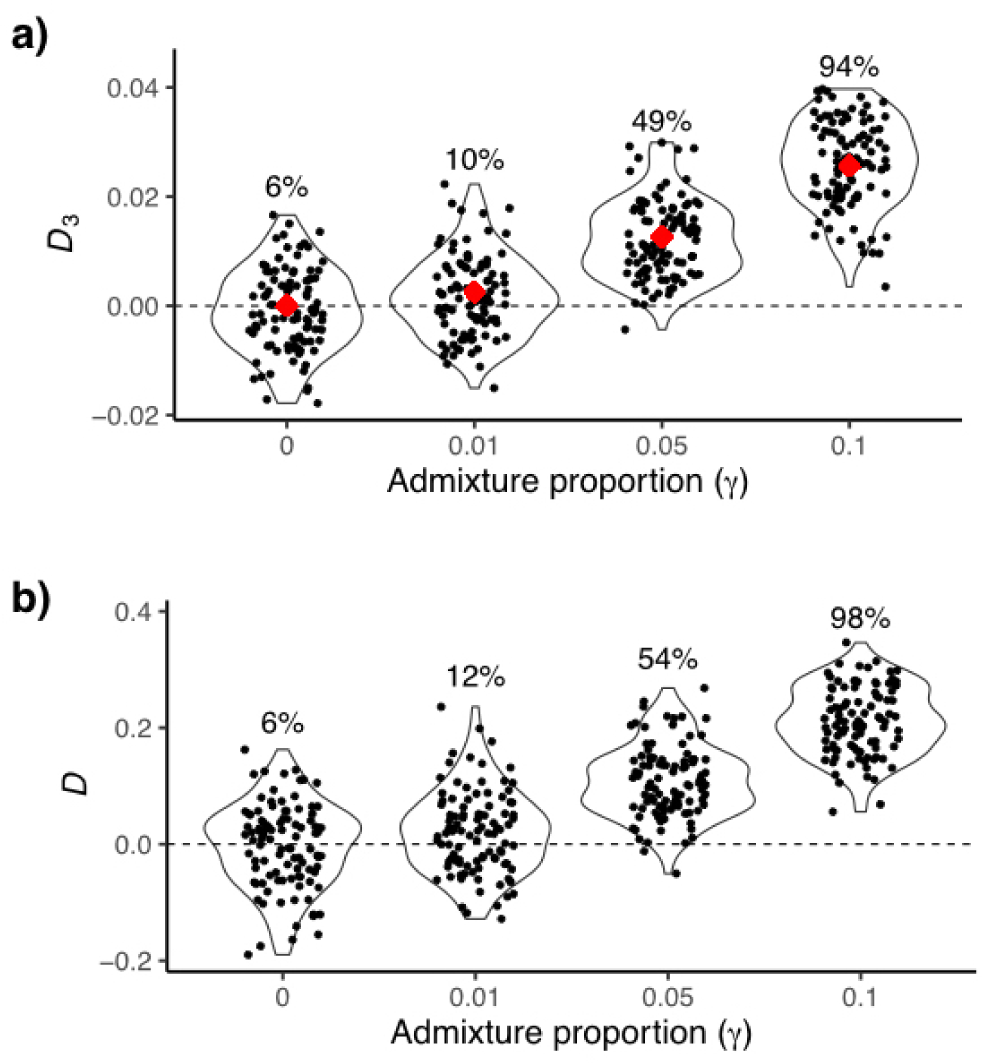
Statistical power of *D*_3_ and *D*. Data were simulated for different proportions of the genome affected by introgression (equivalent to the admixture proportion, *γ*), and significance of each dataset was determined by block bootstrap (see Materials and Methods). Each black point represents the value of either a) *D*_3_ or b) *D* for each simulated dataset, with some horizontal jitter added for clarity. Violin plots are used to display the density of values, and in the top panel the red diamonds represent the expected values of *D*_3_ for different values of *γ*. Percentages reported above each violin plot represent the proportion of simulated datasets that were significantly different from 0 at *P* < 0.05.

In order to directly compare these power calculations to the traditional *D*-test, we included an outgroup in the same simulated datasets (the outgroup was simply ignored for *D*_3_ calculations). As shown in Figure 3b, *D* has only slightly more statistical power, despite requiring more data than *D*_3_. Our results match similar calculations for *D* carried out previously (e.g. Good et al. 2015; Martin et al. 2015), demonstrating the general power of this class of tests to detect introgression between non-sister lineages.

*D*_3_ also has some obvious advantages over similar tests, as it does not require an outgroup (as does the *D*-test) or population samples from three taxa (as does the *f*_3_-test; Reich et al. 2009). Even when data from outgroups are available, if there is either ILS or introgression involving these species the *D*-test may not be appropriate. *D*_3_ can also detect introgression in both directions (i.e. from *B* into *C* and from *C* into *B*), similar to *D* but unlike *f*_3_, which can only detect it in one direction (Peter 2016).

The *D*-test has been used with ancient DNA samples, as in the use of Neandertal sequences in the paper introducing this statistic (Green et al. 2010). Although the expectations of branch lengths for *D*_3_ given here obviously assume that all sequences are sampled from the present (or are sampled contemporaneously from the past), all of the symmetry expectations hold if the ancient sample is the unpaired lineage (i.e. species *C* in Figure 1). Therefore, there may also be limited cases in which *D*_3_ can be applied to ancient samples.

### Assumptions of *D*_3_

Several points about the test introduced here merit further discussion and explanation. Although the expectations underlying *D*_3_ require few assumptions, there are a few things to be cautious about. First, we have assumed that the pairwise distances used as input to *D*_3_ accurately reflect coalescence times. This will only strictly be true for sequences evolving under an infinite sites model with the same shared mutation rate across lineages. Such conditions likely hold only for relatively closely related species, limiting the use of *D*_3_ to recent divergences.

Second, while values of *D*_3_ significantly different from zero can be interpreted as rejecting an ILS-only model (given the above assumptions), such results do not strictly mean that introgression is the cause of rejection. As with the *D*-test, population structure in the ancestor of all three lineages can produce deviations from the ILS-only expectations (Slatkin and Pollack 2008; Durand et al. 2011). In these cases additional analyses may be needed to distinguish among alternative causes of significant *D*_3_ values (e.g. Lohse and Frantz 2014).

Finally, we have assumed here that the rooted species tree is known, even though the test does not require an outgroup. Of course it is often the case that the species tree can be inferred from either smaller amounts of sequence data or morphological characters, and so the species tree may be known despite the lack of genome-scale data from an outgroup taxon. However, if the species relationships are not known, a conservative approach would be to test all three combinations of pairwise distances (i.e. *d*_*B−C*_*–d*_*A−C*_, *d*_*B−C*_*–d*_*A−B*_, and *d*_*A−C*_*–d*_*A−B*_). If all three are significantly different from zero, then it is likely that introgression has acted in the system.

## Materials and Methods

In order to determine the statistical power of the tests discussed here, we simulated multi-locus datasets. For each of four different values of the admixture proportion (γ), we simulated 100 datasets consisting of 1000 non-recombining loci each using the coalescent simulator *ms* (Hudson 2002). The species tree used for all conditions had *t*_1_ = 0.3 and *t*_2_ = 0.6, and simulations with introgression had *t*_*m*_ = 0.05 (in units of 4*N* generations). All simulations also included an outgroup taxon that diverged at *t*_*o*_=4, though data from the outgroup was only used for calculations involving *D*. Gene trees from *ms* were passed to Seq-Gen (Rambaut and Grassly 1997) to simulate 1-kb alignments under the Jukes-Cantor model with *θ*=0.01. All simulation commands are provided in the Appendix.

The resulting datasets of 1000 loci were concatenated together to calculate either *D* or *D*_3_. Significance of each simulated dataset was determined by block bootrapping 1000 times (with block size equal to 10-kb). The resulting values of *D* or *D*_3_ were used to generate a *z* distribution, and a nominal value of *P* < 0.05 was used as a threshold for significance.

## Acknowledgments

The authors are grateful for Mathieu Rouard’s assistance with the *Musa* data, and for discussions with Jeff Good and Eric Stone that helped to improve the manuscript. This work was supported by the Precision Health Initiative of Indiana University and National Science Foundation grant DBI-1564611.

## Appendix

### Model with introgression

When there is introgression, some loci have a history that takes a different path through the species network. For the simplest case with one introgression event, there is one reticulation and therefore one additional “parent tree” embedded in the species network (see Hibbins and Hahn 2019 for full explication). Here we describe the expectations for introgression from species *C* into species *B* (as in Figure 2A in the main text). Other introgression events follow the same logic as this one. The additional parent tree generated by this introgression has lineages *C* and *B* sister to one another, and is defined by two split times: *t*_*m*_ and *t*_2_. The time *t*_2_ is the same as in the species tree, but now lineages *B* and *C* can coalesce starting at time *t*_*m*_ (Figure 2A).

This parent tree can also produce all three possible topologies. Because the *BC* topology is now the one that matches the parent tree, there are two histories with this topology; we denote these *BC*1_2_ and *BC*2_2_. The two topologies discordant with the parent tree from an introgression history are denoted *AB*_2_ and *AC*_2_. The expected frequencies of these topologies are:

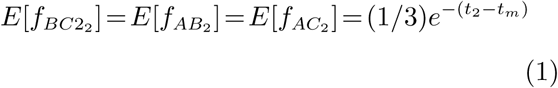

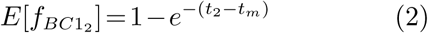

Our goal here is to find the expectation for *D*_3_ (as given in equation 8 in the main text) in the presence of introgression. We therefore require the expected coalescence times *t*_*B−C*_ and *t*_*A−C*_ for each topology from the second parent tree:

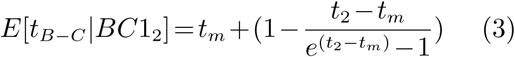

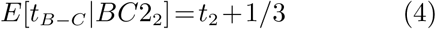

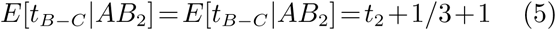

and

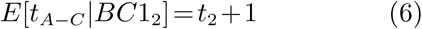

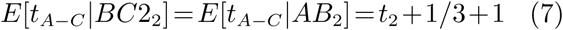

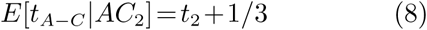

The expected distances between lineages across all loci will be comprised of the average distance across trees with both introgressed and non-introgressed histories. Therefore, we must weight the contributions of each history by the admixture proportion, *γ*, which describes the fraction of the genome following the introgression history (with 1-*γ* following the species history). Combining results on the expected time to coalescence for the species history (given in the main text, and denoted with the subscript “1” here) with the expected times for the introgression history (supplementary equations 1–8), we have:

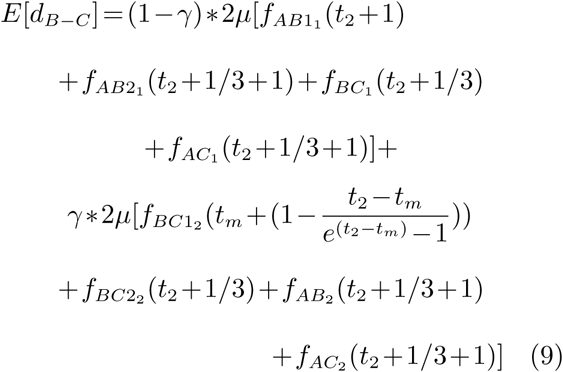

and

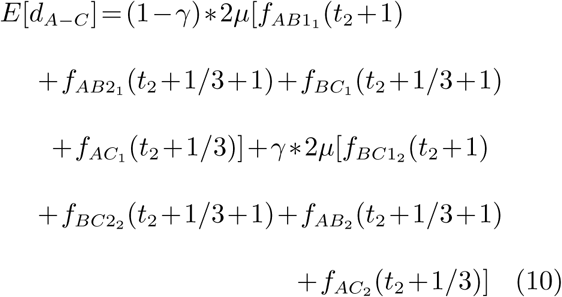

These expected values can be used to find the value of *D*_3_ for any amount of introgression at any time in the past (see, for example, Figure 3 in the main text).

### Simulated alignments with introgression

To quantify the power of *D*_3_ and compare it to the *D*-test, we simulated 1-kb alignments of three focal species plus an outgroup, under four different introgression scenarios. For the scenario with no introgression, we used the following call to ms:

ms 4 1000 -T -I 4 1 1 1 1 -ej 4.0 2 1 -ej 0.6 3 2 -ej 0.3 4 3

For the three scenarios with introgression, we used:

ms 4 1000 -T -I 4 1 1 1 1 -ej 4.0 2 1 -ej 0.6 3 2 -ej 0.3 4 3 -es 0.05 3 tbs -ej 0.05 5 2

We passed values of 0.99, 0.95, and 0.9 to “tbs”, corresponding to admixture proportions of 0.01, 0.05, and 0.1, respectively. All gene trees simulated in ms were passed to Seq-Gen to generate alignments with the following:

seq-gen -m HKY -l 1000 -s 0.01

These alignments were then concatenated and used to estimate the statistics.

## References

Durand, E. Y., N. Patterson, D. Reich, and M. Slatkin. 2011. Testing for ancient admixture between closely related populations. Molecular Biology and Evolution 28:2239–2252.

Edelman, N. B., P. Frandsen, M. Miyagi et al. 2018. Genomic architecture and introgression shape a butterfly radiation. bioRxiv:466292.

Elworth, R. A. L., C. Allen, T. Benedict, P. Dulworth, and L. Nakhleh. 2018. *DGEN*: A test statistic for detection of general introgression scenarios. bioRxiv:348649.

Good, J. M., D. Vanderpool, S. Keeble, and K. Bi. 2015. Negligible nuclear introgression despite complete mitochondrial capture between two species of chipmunks. Evolution 69:1961–1972.

Green, R. E., J. Krause, A. W. Briggs et al. 2010. A draft sequence of the Neandertal genome. Science 328:710–722.

Hibbins, M. S., and M. W. Hahn. 2019. The timing and direction of introgression under the multispecies network coalescent. Genetics 211:1059–1073.

Hudson, R. R. 1983. Testing the constant-rate neutral allele model with protein sequence data. Evolution 37:203–217.

Hudson, R. R. 2002. Generating samples under a Wright–Fisher neutral model of genetic variation. Bioinformatics 18:337338.

Huson, D. H., T. Klopper, P. J. Lockhart, and M. A. Steel. 2005. Reconstruction of reticulate networks from gene trees. Research in Computational Molecular Biology, Proceedings 3500:233–249.

Lohse, K., and L. A. Frantz. 2014. Neandertal admixture in Eurasia confirmed by maximum-likelihood analysis of three genomes. Genetics 196:1241–1251.

Mallet, J., N. Besansky, and M. W. Hahn. 2016. How reticulated are species? Bioessays 38:140–149.

Martin, S. H., J. W. Davey, and C. D. Jiggins. 2015. Evaluating the use of ABBA-BABA statistics to locate introgressed loci. Molecular Biology and Evolution 32:244–257.

Meng, C., and L. S. Kubatko. 2009. Detecting hybrid speciation in the presence of incomplete lineage sorting using gene tree incongruence: a model. Theoretical Population Biology 75:35–45.

Pease, J. B., and M. W. Hahn. 2013. More accurate phylogenies inferred from low-recombination regions in the presence of incomplete lineage sorting. Evolution 67:2376–2384.

Pease, J. B., and M. W. Hahn. 2015. Detection and polarization of introgression in a five-taxon phylogeny. Systematic Biology 64:651–662.

Peter, B. M. 2016. Admixture, population structure, and *F*-statistics. Genetics 202:1485–1501.

Rambaut, A., and N. C. Grassly. 1997. Seq-Gen: an application for the Monte Carlo simulation of DNA sequence evolution along phylogenetic trees. CABIOS 13:235–238.

Reich, D., K. Thangaraj, N. Patterson, A. L. Price, and L. Singh. 2009. Reconstructing Indian population history. Nature 461:489–494.

Rouard, M., G. Droc, G. Martin et al. 2018. Three new genome assemblies support a rapid radiation in *Musa acuminata* (wild banana). Genome Biology and Evolution 10:31293140.

Slatkin, M., and J. L. Pollack. 2008. Subdivision in an ancestral species creates asymmetry in gene trees. Molecular Biology and Evolution 25:2241–2246.

Yu, Y., C. Than, J. H. Degnan, and L. Nakhleh. 2011. Coalescent histories on phylogenetic networks and detection of hybridization despite incomplete lineage sorting. Systematic Biology 60:138–149.

